# Barcode Crosstalk in ONT Multiplex Sequencing: Quantification and Mitigation Strategies

**DOI:** 10.64898/2026.03.27.714689

**Authors:** Sebastian A. Scharf, Philipp Spohr, Max J. Ried, Rainer Haas, Gunnar W. Klau, Birgit Henrich, Klaus Pfeffer

## Abstract

Multiplexing samples in long-read sequencing with Oxford Nanopore Next Generation Sequencing Technology (ONT) by ligating specific native barcodes to individual DNA samples enables significant increases of high throughput sequencing combined with a significant reduction of sequencing costs. However, this advantage carries the risk of barcode misassignment / crosstalk. Employing ONT multiplex sequencing with samples, we observed misassigned barcodes so called barcode crosstalk, after ONT library preparation according to the standard protocol, particularly in samples with low input DNA concentrations. We assumed that these barcode misassignments are largely due to misligation of remaining native barcodes during subsequent the subsequent sequencing adapter ligation.

To systematically investigate and quantify barcode crosstalk, genomic DNA (gDNA) from four bacterial type strains with different DNA input concentrations was prepared using three protocols for library preparation: the Nanopore standard protocol (protocol A: version valid until July 2, 2025) the new Nanopore protocol (protocol B: version from July 2, 2025), and an in house protocol with pooling of the barcoded samples only after the sequencing adapter ligation step (protocol C: in house). All samples were sequenced on a Nanopore PromethIon device.

The results clearly showed that the use of protocol A resulted in a pronounced barcode crosstalk especially detectable in samples with low DNA input concentrations (up to 2.4% misassigned reads). The ONT adjustment in protocol B (altered washing buffer vs. protocol A) significantly alleviated the barcode crosstalk to below 0.01%, whereas protocol C eliminated barcode crosstalk virtually completely.

These observations emphasize that sequencing results obtained with older ONT native barcoding protocol variants should be critically reviewed. The newer ONT barcoding protocol is preferable for sequencing, but it does not completely eliminate the barcode crosstalk effect. In conclusion, for low DNA input and high accuracy sequencing, protocol C is recommended.

## Introduction

ONT long read whole genome sequencing has become one of the main tools for the decryption of the composition of clinical metagenomic samples, such as stool, wound swabs or saliva ^1–4^. Especially the ability to detect eukaryotic organisms, bacteria, DNA viruses, and archaea in a metagenomic sample with a single analysis is still offset by the high cost of Nanopore sequencing compared to traditional 16S or 18S rDNA-based PCR methods. These costs are largely determined by the Nanopore flow cells. Multiplexing can significantly reduce the sequencing costs per sample. To this end, ONT has developed protocols such as the Ligation Sequencing gDNA – Native Barcoding Kit 24 V14, which allows up to 24 samples to be sequenced simultaneously on a single flow cell. This is achieved by ligating a unique native barcode to the DNA fragments of each individual sample, theoretically allowing the sequence data to be uniquely assigned to the corresponding samples. However, barcoding can cause misassignments leading to distortion of quantitative sequencing results, false positives, and aberrations in diversity assessments impairing reproducibility and accuracy of the method.

Sequencing of multiplexed low input DNA metagenomic samples together with DNA derived from single bacterial isolates using the Native Barcoding Kit on one flowcell, we noticed a significant fraction of reads in the metagenomic samples appearing to belong to the bacterial isolates. The standard Nanopore protocol involves two ligation steps: In the first step, native barcodes are ligated to the prepared DNA, followed by a second step in which the sequencing adapters are ligated to the already pooled and barcoded DNA. After isolating the error source by performing additional experiments to exclude direct contamination during sample preparation or lab-resident microbes, we hypothesized that barcode adapters might remain after washing and bind to DNA from other samples in the sample pool during the adapter ligation step.

As reported previously for multiplexed Illumina sequencing ^5^, we show in this study that barcode crosstalk is also a significant but, up to now, vastly underestimated effect in ONT multiplex sequencing ^6–8^ that one should be aware of, especially when only samples with low biomass and / or with low DNA quantities are available which are below the recommended amounts of ONT. We have developed a modified sample preparation protocol that even under these conditions mitigates barcode crosstalk to a minimum thus increasing ONT multiplex sequencing accuracy and reproducibility.

## Results

In order to investigate and to quantify the extent of barcode crosstalk with ONT multiplexed sequencing, we performed three independent sequencing runs, each using three different sample preparation protocols A, B and C (see Fig. 1). The first was the original ONT Ligation sequencing gDNA - Native Barcoding protocol (tSQK-NBD114.24 / NBE_9169_v114_revU_30Jan2025), where native barcode ligation samples are pooled after washing and washing is performed with 80% ethanol. Protocol B is an advanced variant of protocol A, which was published by ONT (SQK-NBD114.24 / NBE_9169_v114_revV_02Jul2025) to mitigate barcode crosstalk. Instead of ethanol, here short fragment buffer (SFB) is used for washing after barcode ligation. The third protocol C (BEAP) is our developed workflow, using the tSQK-NBD114.24 / NBE_9169_v114_revU_30Jan2025 reagents protocol with the important modification that individual samples are pooled only at the latest timepoint after the sequence adapter ligation step to virtually exclude incorrect barcode ligations.

**Fig. 1:**
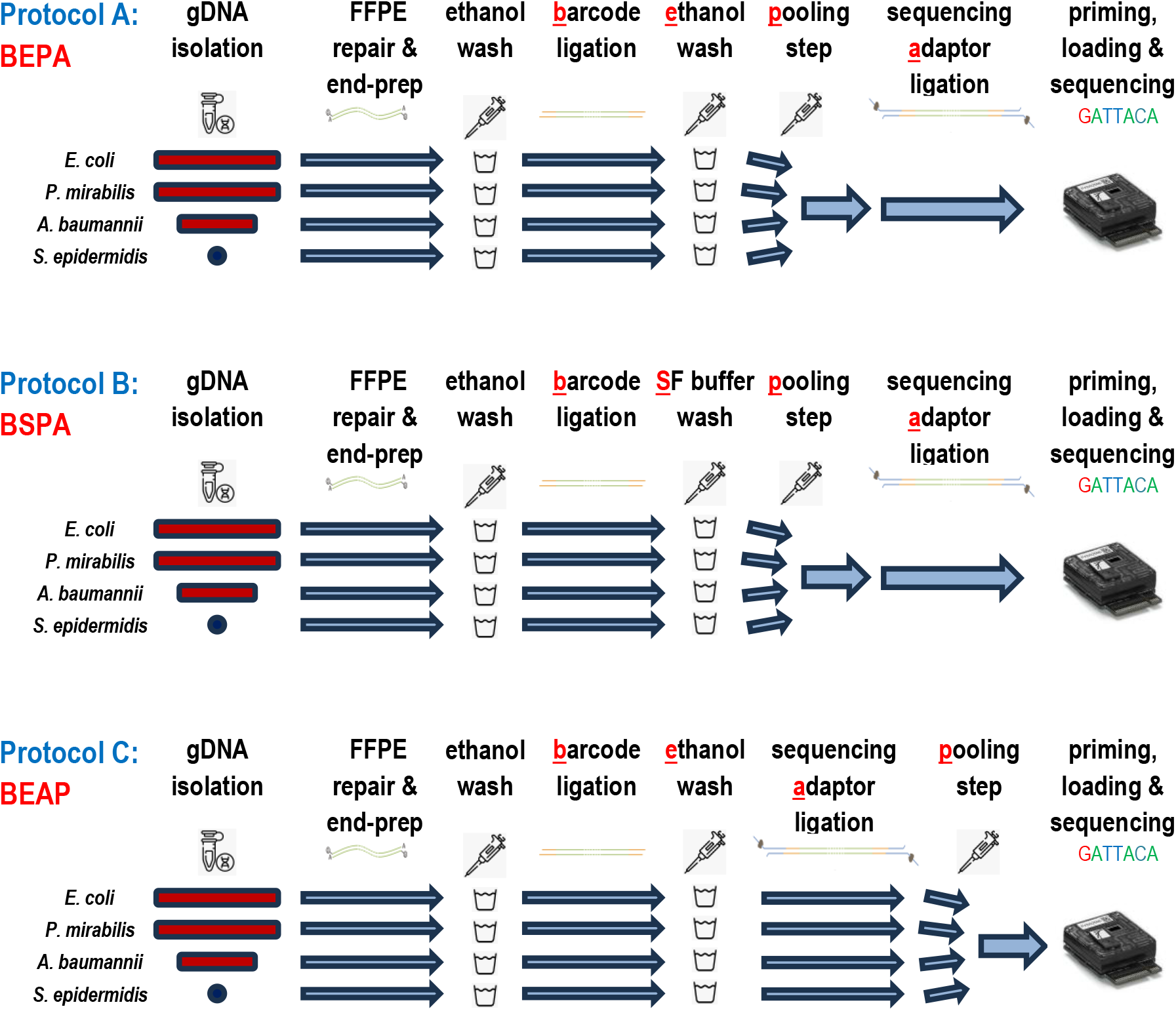
Protocol schemes. Workflows of the three sample preparation protocols for NGS sequencing used in this study. Protocol A (BEPA) was the SQK-NBD114.24 Nanopore Protocol in the version NBE_9169_v114_revU_30Jan2025, where samples are pooled after the barcode ligation and washed with ethanol. Protocol B (BSPA) is an advanced variant of protocol A, which was published by Oxford Nanopore in July 2025 (NBE_9169_v114_revV_02Jul2025) to mitigate barcode crosstalk. Instead of ethanol, short fragment buffer (SFB) is used for washing after the barcode ligation step. The last protocol C (BEAP) is our own variant, where samples are pooled at the latest timepoint after the sequence adapter ligation step to virtually exclude incorrect barcode ligations.

To quantify the extent of barcode crosstalk among sequencing samples with varying DNA input concentrations, the three sample preparation protocol variants were performed with gDNA from four different bacterial type strains (*Escherichia coli* (ATCC 35218), *Proteus mirabilis, Acinetobacter baumannii, S. epidermidis*) on four different dilution levels: 1:1 (corresponding to 400 ng genomic DNA in 12 µl Tris-Buffer as recommended by ONT) for *E. coli* and in dilutions of 1:10 (40 ng gDNA) for *P. mirabilis* (ATCC 29906), 1:100 (4 ng gDNA) for *A. baumannii* (ATCC BAA 747), and 1:1000 (0.4 ng gDNA) for *S. epidermidis* (ATCC 12228) in TRIS-buffer. This experimental design allowed the barcode crosstalk effect to be systematically quantified as a function of the respective gDNA input quantity.

After performing the experiments (n=3) we observed an expected approx. logarithmic decrease in the total number of reads with 10-fold diluted concentrations of input DNA (Table 1). Protocol A, the Nanopore variant with ethanol as wash buffer and protocol B, where in the first washing step, ethanol was replaced by SFB, showed similar readcount losses. Compared to the total readcounts of the 1:1 diluted samples (Protocol A: 5.42x10^6^ (+/-1.58x10^6^); Protocol B: 4.66x10^6^ (+/-2.52 x10^6^)) the decrease of reads was about 10-23 fold for the DNA dilution of 1:10, 130-190 fold for 1:100 and 2500-7500 fold for 1:1000. Protocol C, with the pooling step after adapter ligation, led to lower read counts than the other protocols (1:1 dilution: 4.58x10^6^ (+/-2.30x10^6^)) with a higher readcount loss of 30 fold (1:10), 470 fold (1:100) and 11739 fold (1:1000), respectively.

**Table 1.** Number of reads (mean values) per type strain after read mapping with minimap2. After carrying out sample preparation according to the three protocols (see Fig.1) sequencing the multiplexed samples was performed on a ONT PromethION platform. Basecalling was done with Dorado vers. 7.9.8 using the high-accuracy model. Reads that could be mapped to one of the reference genomes (excluding masked regions in the reference genomes or reads that could not be mapped) were counted and are depicted. The experiment was repeated independently three times.

| Protocol | Barcode used for ligation of gDNA isolated from | Sequence identified originating from [mean values] |  |  |  |
| --- | --- | --- | --- | --- | --- |
|  |  | *E. coli* | *P. mirabilis* | *A. baumannii* | *S. epidermidis* |
| A [BEPA] | *E. coli* | 5,420,544.33 | 3.33 | 0.33 | 0.00 |
|  | *P. mirabilis* | 11.67 | 263,683.33 | 0.00 | 0.00 |
|  | *A. baumannii* | 11.00 | 0.00 | 29,645.00 | 0.00 |
|  | *S. epidermidis* | 21.00 | 0.00 | 0.00 | 747.33 |
| B [BSPA] | *E. coli* | 4,656,477.33 | 1.67 | 0.33 | 0.33 |
|  | *P. mirabilis* | 0.33 | 212,586.67 | 0.00 | 0.00 |
|  | *A. baumannii* | 0.33 | 0.00 | 27,790.00 | 0.00 |
|  | *S. epidermidis* | 0.00 | 0.00 | 0.00 | 457.00 |
| C [BEAP] | *E. coli* | 4,584,124.33 | 2.33 | 0.00 | 0.00 |
|  | *P. mirabilis* | 0.00 | 111,720.67 | 0.00 | 0.00 |
|  | *A. baumannii* | 0.00 | 0.00 | 9,202.00 | 0.00 |
|  | *S. epidermidis* | 0.00 | 0.00 | 0.00 | 236.33 |

Using protocol A [BEPA], a clear barcode crosstalk effect became apparent, especially in the samples with lower amounts of input DNA (Table 2). The samples incorrectly gained reads that had the expected barcode for this sample, but the barcode was found ligated to gDNA from other type strains - i.e., expected barcode, but incorrect DNA.

**Table 2.** Barcode crosstalk - gained incorrect reads (%) means ± standard deviations. By changing the washing buffer from ethanol to SFB, according to the updated Nanopore protocol (protocol B [BSPA]), barcode crosstalk was drastically reduced: 0.0001% (+/-0.0000%) incorrect reads (1:1 dilution), 0.0001% (+/-0.0002%) (1:10 dilution), 0.0013% (+/-0.0019%) (1:100 dilution) and 0.0000% (+/-0.0000%) (1:1000 dilution), however, still detactable.

| Protocol | Barcode used for ligation<br>of gDNA isolated from | Input gDNA<br>[ng] | % incorrect<br>reads | ± | SD |
| --- | --- | --- | --- | --- | --- |
| **A [BEPA]** | *E. coli* | 400.0 | 0.0001 | ± | 0.0000 |
|  | *P. mirabilis* | 40.0 | 0.0042 | ± | 0.0028 |
|  | *A. baumannii* | 4.0 | 0.0357 | ± | 0.0242 |
|  | *S. epidermidis* | 0.4 | 2.3891 | ± | 0.8920 |
| **B [BSPA]** | *E. coli* | 400.0 | 0.0001 | ± | 0.0000 |
|  | *P. mirabilis* | 40.0 | 0.0001 | ± | 0.0002 |
|  | *A. baumannii* | 4.0 | 0.0013 | ± | 0.0023 |
|  | *S. epidermidis* | 0.4 | 0.0000 | ± | 0.0000 |
| **C [BEAP]** | *E. coli* | 400.0 | 0.0000 | ± | 0.0000 |
|  | *P. mirabilis* | 40.0 | 0.0000 | ± | 0.0000 |
|  | *A. baumannii* | 4.0 | 0.0000 | ± | 0.0000 |
|  | *S. epidermidis* | 0.4 | 0.0000 | ± | 0.0000 |

The percentage of incorrect assigned reads increased from 0.0001% (+/-0.0000%) (1:1 dilution) over 0.0042% (+/- 0.0023%) (1:10 dilution) to 0.0357% (+/- 0.0198%). At a dilution of 1:1000, only about 97.6% (+/- 0.72%) of the reads could be classified correctly.

Clear improvement was only achieved by adjusting the workflow of the protocol by combining the samples after sequence adapter ligation (protocol C [BEAP]). Here, in total, only seven reads were incorrect for all DNA input / dilution levels, resulting in an average percentage of incorrect reads below 3,9E-05.

In addition to incorrectly gained reads, misassigned barcodes also cause samples to lose correct reads to other samples - i.e., incorrect barcode, but correct DNA. in this analysis of the sequencing data also protocol C [BEAP] had the lowest percentage of incorrect barcodes, i.e., incorrect binning (Table 3).

**Table 3.**
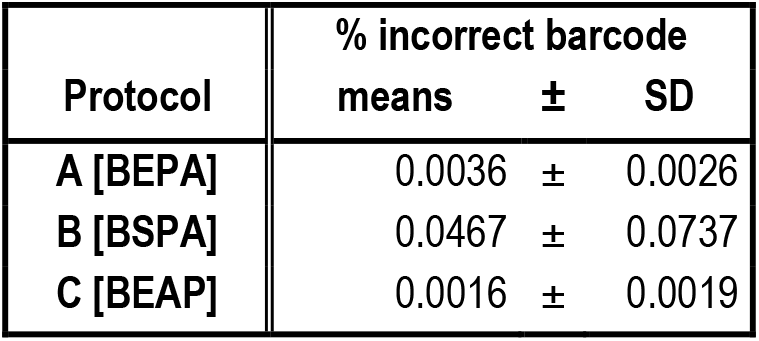
Barcode crosstalk - lost correct reads (%) means ± standard deviations. Using protocol A, the samples prepared lost more reads as in protocol C [BEAP] (Table 1, Table S2). Surprisingly, in protocol B, however, the highest amounts of gDNA were ligated to incorrect barcodes (Table 3), arguing that the modified ONT protocol with SFB wash buffer still leaves high amounts of barcodes in the reaction mix. Thus, the extent of barcode crosstalk is considerable in samples with low concentrations: incorrect barcoding of gDNA can significantly distort the composition of the sample by ligating incorrect barcodes to gDNA thus influencing accuracy and reproducibility of sequencing results.

| Protocol | % incorrect barcode |  |  |
| --- | --- | --- | --- |
|  | means | ± | SD |
| A [BEPA] | 0.0036 | ± | 0.0026 |
| B [BSPA] | 0.0467 | ± | 0.0737 |
| C [BEAP] | 0.0016 | ± | 0.0019 |

Taken together, in our investigation, it becomes obvious that barcode crosstalk mainly can be detected from the high DNA input sample into the less concentrated samples (see Table 1), possibly due to an “residual” amount of free barcodes in the diluted samples and non-barcode ligated DNA fragments in the high DNA input samples after the barcode ligation step. In percentage terms, false reads therefore occurred mainly in the samples with lower DNA input. This barcode crosstalk problem can be mitigated by the adjustment of the Nanopore workflow protocol now using SF buffer instead of ethanol for the washing step after barcode ligation. At a dilution of 1:1000, we observed virtually no barcode crosstalk in our experiments with protocol B [BSPA]. Importantly, the protocol we developed (protocol C [BEAP], where samples were only pooled after sequencing adapter ligation, virtually abrogated barcode crosstalk. However, this method is more cost-intensive than the adjustment proposed by Nanopore, which only requires a modified wash buffer. In the case of the necessity of highest accuracy, protocol C with pooling of the samples only after all ligation steps should be preferred.

## Discussion

While barcode crosstalk is a known phenomenon in multiplexed Illumina sequencing ^6^ it is so far surprisingly underreported in Nanopore sequencing and rarely quantified ^7^ . Xu et al. identified 0.056% of total reads in their sequencing data being wrongly assigned to incorrect barcodes. A note in the paper of Wick and coworkers discusses the effect briefly and estimates it at 0.3% of all reads being misassigned ^9^. Recently Dai et al. published a preprint discussing the effect and quantified it through an experimental setup as well. Here, the estimated percentage of misassigned reads was reported at 0.36% ^10^. ONT is also aware of barcode crosstalk and published a revised version of its sequencing protocol (2.7.2025). According to the manufacturer, the washing buffer after barcode ligation was adjusted from ethanol to SFB washing buffer in this new version of the protocol, specifically to minimize barcode crosstalk ^11^. Since multiplexing using barcoding is common practice to save costs, it is crucial to understand the size of the effect. Barcode crosstalk mitigation strategies especially for low input DNA concentrations are essential, since the resulting error rate is unacceptable when analyzing low abundance samples or when low fractions of reads in a sample are required to be informative. We propose a variation of the ONT protocol and compared it to the modified ONT protocol also addressing the barcode crosstalk issue that was released 2.7.2025.

Our results indicate that barcode crosstalk occurs during the second ligation step of the ONT protocols, causing excess barcodes from the pool to be ligated nonspecifically to up to now non-barcode ligated, gDNA from other samples, since physical purification steps and chemical ligation reactions do not have 100% efficiency due to thermodynamic constrains. Oxford Nanopore Technologies is aware of the barcode crosstalk effect and adjusted the protocol so that SFB buffer is used for washing after barcode ligation instead of ethanol. Due to the use of the SFB buffer, excess barcodes were still found ligated to gDNA from different origin, but were found less frequently, this resulting in a significant reduction in barcode crosstalk.

This modified ONT protocol variant [BSPA] is significantly more cost-effective and less time-consuming than newly established protocol C [BEAP] that we developed in which the samples are only pooled after sequence adapter ligation. While the updated Nanopore protocol only requires more SFB buffer—currently available as an add-on to the kit, protocol C significantly increases the required amounts of enzymes and buffer solutions and is more labor-intensive. This is because the samples must be processed individually rather than collectively after barcoding. Instead of up to 144 samples (24 barcodes × 6 pool reactions), only six samples can be processed with one sequencing kit, which significantly increases the cost per sample. Another disadvantage of protocol C [BEAP] is the increased time required, as each sample must be processed separately rather than as a common pool. However, it must be taken into account that switching to SFB buffers only reduces the problem, but does not solve it completely, which can only be achieved by later pooling.

While the barcode crosstalk effect itself is evident with our results, our experiments are limited by resolution. A higher sequencing depth could quantify the effect with even more confidence and precision. We also believe that the binding of adapters could vary slightly between different barcodes, kits and DNA content. Additional experiments should quantify the effects of those factors. The barcode misligation problem has already been discussed in another study ^10^. There, too, it was shown that modifications to the Nanopore protocol—in particular, the later pooling of samples, as we did in protocol variant C, led to a significant reduction in barcode crosstalk. In comparison to Dai et al., who used DNA from *P. vulgatus, M. hominis* and *C. striatum*, we used DNA from *E. coli, P. mirabilis, A. baumanii* and *S. epidermidis* type strains. With this approach. we demonstrate that the misligation effect is independent of the bacterial species sequenced. While the concentration of the DNA they used remained the same, we were able to show with our dilution series that the crosstalk effect increases as the concentration of the input DNA decreases.

The barcode crosstalk effect was measured quantitatively using the tested dilution series. This revealed a significant increase in the effect as the input DNA concentration decreased, resulting in up to 2.4% of reads being misclassified at the lowest dilution level with the old Nanopore protocol. Against this background, ONT nanopore sequenced samples with low abundance that were performed according to the old protocol variant should be critically reviewed. The change in the washing buffer (protocol B) significantly reduces the crosstalk effect, with only variant C (pooling after adapter ligation) virtually abrogates the barcode crosstalk. It should therefore be considered especially for samples with low input, i.e. when analyzing cell-free DNA from blood samples or for samples with low abundance gDNA content such as microbiomes from spinal fluid, urine etc. as well as low biomass samples where cross-sample contamination is of concern.

## Conclusion

Barcode crosstalk is a relevant but previously underestimated effect in sequencing with ONT technology, especially in samples with low biomass or with DNA inputs below the recommended amount. Our quantitative analyses with dilution series showed that the crosstalk effect increases significantly as DNA concentration decreases and can lead to misclassification of up to 2.4% of reads when using the old protocol. The new Nanopore protocol significantly reduces this effect, while only our protocol variant, which uses pooling after the adapter ligation, nearly eliminates the crosstalk effect entirely.

## Material and Methods

The following ATTC type strains were used for our experiments: *Escherichia coli* ATCC 35218, *Proteus mirabilis* ATCC 29906, *Acinetobacter baumannii* ATCC BAA747, *Staphylococcus epidermidis* ATCC 12228.

The strains were cultured on COS agar plates (Columbia agar + 5% sheep blood; Biomerieux, Marcy-l’Étoile, France) and the species identity was confirmed by MALDI-TOF (Vitek MS Prime; Biomerieux, Marcy-l’Étoile, France). One colony of each isolate was taken and cultivated in 1.8 mL LB liquid media. After 12 h of incubation at 37°C the suspension was taken and genomic DNA was extracted with the DNeasy Ultra Clean Microbial Kit ((Qiagen, Hilden, Germany) according to the protocol of the manufacturer and stored at 4°C. The concentration of the extracted DNA was determined by fluorometry using the Invitrogen Qubit 4 Fluorometer (Thermo Fisher Scientific).

Whole genome sequencing of the isolates was performed using the Native Barcoding Kit 24 V14 (SQK-NBD114.24; Oxford Nanopore Technologies, Oxford, UK). For library preparation three different protocols were performed (see Fig. 1). Protocol A (SQK-NBD114.24 Nanopore Protocol in the version NBE_9169_v114_revU_30Jan2025), Protocol B (SQK-NBD114.24 Nanopore Protocol in the version NBE_9169_v114_revV_02Jul2025) and protocol C (our own variant,based on SQK-NBD114.24 Nanopore Protocol in the version NBE_9169_v114_revU_30Jan2025 where samples are pooled at a later timepoint after the adapter ligation).

The DNA libraries were loaded on PromethION R10.4.1 flow cells and sequenced on the PromethION PC24 with the MinKNOW v24.06.10 software. After sequencing, raw data were base-called using Dorado 7.9.8 using the high-accuracy model. The experiment was performed three times, means and standard deviations were calculated from this data.

A minimap2 database was constructed by using the ATCC sequence references (*Escherichia coli* (ATCC 35218), *Proteus mirabilis* (ATCC 29906), *Acinetobacter baumannii* (ATCC BAA 747) and *Staphylococcus epidermidis* (ATCC 12228) and performing a pairwise alignment with nucmer 3.23 and a subsequent identification of homologous regions, which were masked and added as separate, additional references. The references did not contain any of the SQK-NBD114.24 barcode sequences. Reads were mapped using minimap2 and only counted if a) they uniquely mapped to a reference, b) the alignment spanned at least 2.000 bases in the query (implicitly imposing a read-length requirement), c) at least 70% of the query mapped to the reference with d) less than 5% non-matches, e) achieving a mapping quality of 60. In addition, we required reads to have the correct barcode sequence present without any sequencing error at both ends of a read. As a result, we discarded a large fraction of informative reads but were able to achieve a high level of confidence regarding the analyzed ones (see Table S1, unmapped reads).

## Data and code availability

The analysis code can be found at https://github.com/AlBi-HHU/barcode_crosstalk_examination. The sequencing runs are available at the ENA under the study accession PRJEB105337; since ENA currently does not accept pod5 format we only uploaded the HAC fastq files, the pod5 files are locally archived and can be made available on request. We plan on uploading them as soon as pod5 is supported.

## Author contribution statement

SeS and PS conceptualized the project. SeS conducted the wetlab experiments. PS performed the bioinformatics analysis. MR provided technical support. SeS, PS, and MR developed wetlab mitigation strategies. RH, GK, BH, KP supervised the project. All authors have (proof-)read and agreed to the final submission.

## Acknowledgments

The study was funded and supported by the Jürgen Manchot Foundation. Computational support and infrastructure were provided by the “Centre for Information and Media Technology” (ZIM) at the University of Düsseldorf (Germany). We thank Alexander Dilthey and Anna Rommerskirchen for valuable and constructive discussions. Technical support regarding sequencing hardware were provided by the Genomics and Transcriptomics Laboratory (GTL) of the Biological and Medical Research Centre (BMFZ), Medical Faculty, Heinrich-Heine-University Düsseldorf, Düsseldorf, Germany.

## Supplementary Information

**Table S1.**
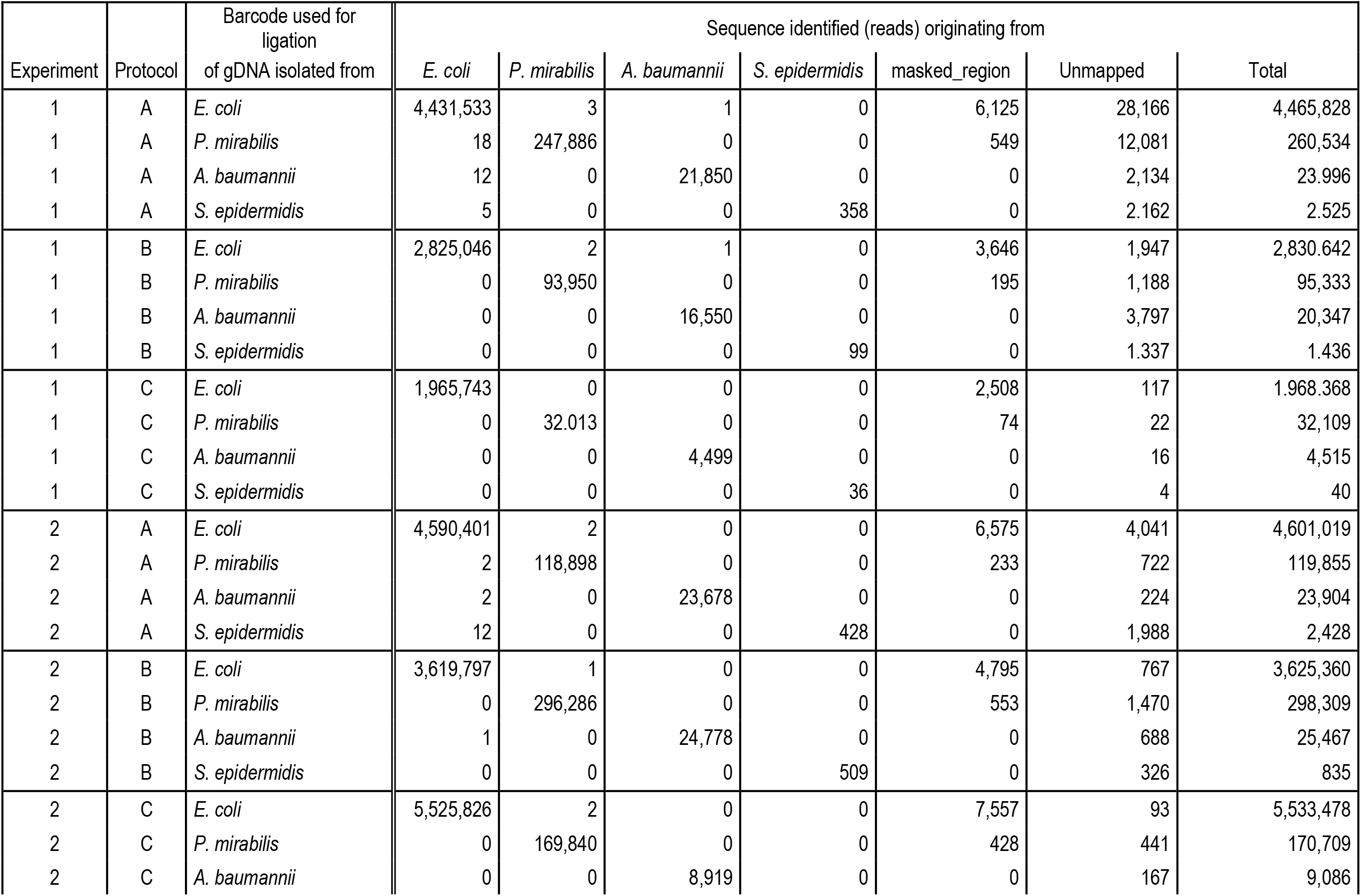
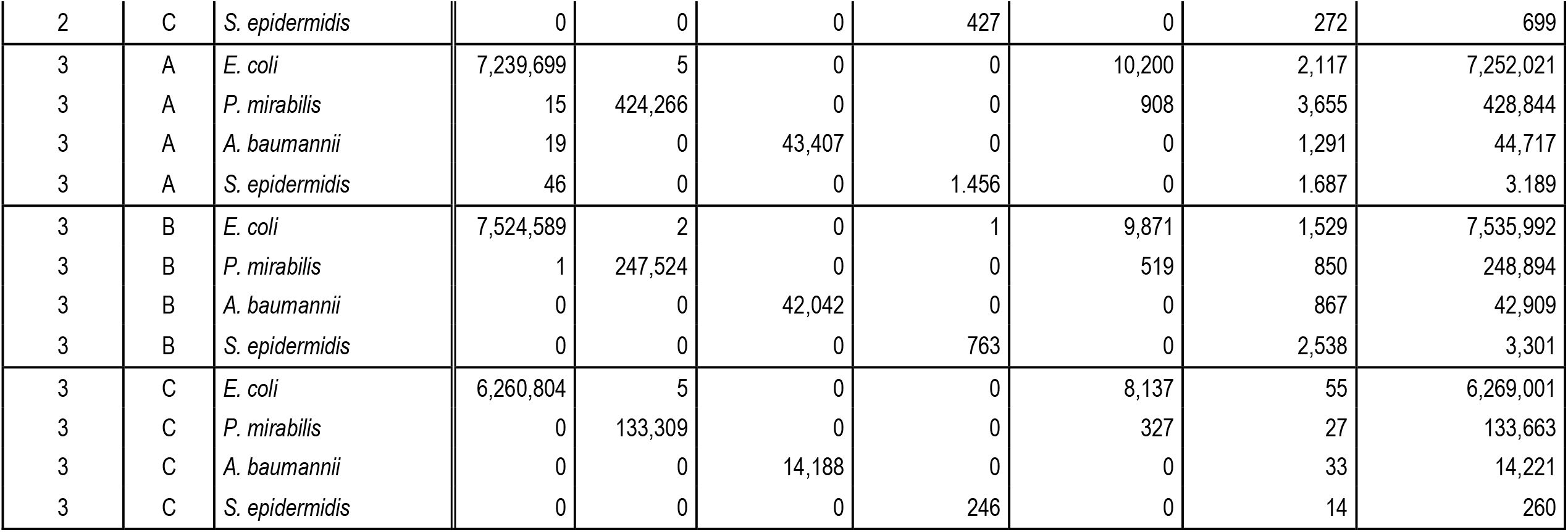
Sequencing results from 3 independent experiments using the three different sample preparation protocols. For further information please refer to legend of Fig. 1.

## Notes

https://github.com/AlBi-HHU/barcode_crosstalk_examination

https://www.ebi.ac.uk/ena/browser/view/PRJEB105337

### Competing Interest Statement

The authors have declared no competing interest.

