## Supplementary Table 1 for "Barcode Crosstalk in ONT Multiplex Sequencing: Quantification and Mitigation Strategies"

### Supplementary Information

| Experiment | Protocol | Barcode used for ligation of gDNA isolated from | Sequence identified (reads) originating from | | | | | | |
|---|---|---|---|---|---|---|---|---|---|
| | | | *E. coli* | *P. mirabilis* | *A. baumannii* | *S. epidermidis* | masked_region | Unmapped | Total |
| 1 | A | *E. coli* | 4,431,533 | 3 | 1 | 0 | 6,125 | 28,166 | 4,465,828 |
| 1 | A | *P. mirabilis* | 18 | 247,886 | 0 | 0 | 549 | 12,081 | 260,534 |
| 1 | A | *A. baumannii* | 12 | 0 | 21,850 | 0 | 0 | 2,134 | 23.996 |
| 1 | A | *S. epidermidis* | 5 | 0 | 0 | 358 | 0 | 2.162 | 2.525 |
| 1 | B | *E. coli* | 2,825,046 | 2 | 1 | 0 | 3,646 | 1,947 | 2,830.642 |
| 1 | B | *P. mirabilis* | 0 | 93,950 | 0 | 0 | 195 | 1,188 | 95,333 |
| 1 | B | *A. baumannii* | 0 | 0 | 16,550 | 0 | 0 | 3,797 | 20,347 |
| 1 | B | *S. epidermidis* | 0 | 0 | 0 | 99 | 0 | 1.337 | 1.436 |
| 1 | C | *E. coli* | 1,965,743 | 0 | 0 | 0 | 2,508 | 117 | 1.968.368 |
| 1 | C | *P. mirabilis* | 0 | 32.013 | 0 | 0 | 74 | 22 | 32,109 |
| 1 | C | *A. baumannii* | 0 | 0 | 4,499 | 0 | 0 | 16 | 4,515 |
| 1 | C | *S. epidermidis* | 0 | 0 | 0 | 36 | 0 | 4 | 40 |
| 2 | A | *E. coli* | 4,590,401 | 2 | 0 | 0 | 6,575 | 4,041 | 4,601,019 |
| 2 | A | *P. mirabilis* | 2 | 118,898 | 0 | 0 | 233 | 722 | 119,855 |
| 2 | A | *A. baumannii* | 2 | 0 | 23,678 | 0 | 0 | 224 | 23,904 |
| 2 | A | *S. epidermidis* | 12 | 0 | 0 | 428 | 0 | 1,988 | 2,428 |
| 2 | B | *E. coli* | 3,619,797 | 1 | 0 | 0 | 4,795 | 767 | 3,625,360 |
| 2 | B | *P. mirabilis* | 0 | 296,286 | 0 | 0 | 553 | 1,470 | 298,309 |
| 2 | B | *A. baumannii* | 1 | 0 | 24,778 | 0 | 0 | 688 | 25,467 |
| 2 | B | *S. epidermidis* | 0 | 0 | 0 | 509 | 0 | 326 | 835 |
| 2 | C | *E. coli* | 5,525,826 | 2 | 0 | 0 | 7,557 | 93 | 5,533,478 |
| 2 | C | *P. mirabilis* | 0 | 169,840 | 0 | 0 | 428 | 441 | 170,709 |
| 2 | C | *A. baumannii* | 0 | 0 | 8,919 | 0 | 0 | 167 | 9,086 |

| | | | | | | | | | |
|---|---|---|---|---|---|---|---|---|---|
| 2 | C | *S. epidermidis* | 0 | 0 | 0 | 427 | 0 | 272 | 699 |
| 3 | A | *E. coli* | 7,239,699 | 5 | 0 | 0 | 10,200 | 2,117 | 7,252,021 |
| 3 | A | *P. mirabilis* | 15 | 424,266 | 0 | 0 | 908 | 3,655 | 428,844 |
| 3 | A | *A. baumannii* | 19 | 0 | 43,407 | 0 | 0 | 1,291 | 44,717 |
| 3 | A | *S. epidermidis* | 46 | 0 | 0 | 1.456 | 0 | 1.687 | 3.189 |
| 3 | B | *E. coli* | 7,524,589 | 2 | 0 | 1 | 9,871 | 1,529 | 7,535,992 |
| 3 | B | *P. mirabilis* | 1 | 247,524 | 0 | 0 | 519 | 850 | 248,894 |
| 3 | B | *A. baumannii* | 0 | 0 | 42,042 | 0 | 0 | 867 | 42,909 |
| 3 | B | *S. epidermidis* | 0 | 0 | 0 | 763 | 0 | 2,538 | 3,301 |
| 3 | C | *E. coli* | 6,260,804 | 5 | 0 | 0 | 8,137 | 55 | 6,269,001 |
| 3 | C | *P. mirabilis* | 0 | 133,309 | 0 | 0 | 327 | 27 | 133,663 |
| 3 | C | *A. baumannii* | 0 | 0 | 14,188 | 0 | 0 | 33 | 14,221 |
| 3 | C | *S. epidermidis* | 0 | 0 | 0 | 246 | 0 | 14 | 260 |

**Table S1. Sequencing results from 3 independent experiments using the three different sample preparation protocols**. For further information please refer to legend of Fig. 1.
